# State or trait? MRS-measured GABA and Glutamate concentrations are not modulated by task demand and do not robustly predict task performance

**DOI:** 10.1101/543140

**Authors:** Lotte Talsma, Anouk van Loon, H. Steven Scholte, Heleen A. Slagter

## Abstract

Over the past few years, Magnetic Resonance Spectroscopy (MRS) has become a popular method to non-invasively study the relationship between in-vivo concentrations of neurotransmitters such as GABA and Glutamate and cognitive functions in the human brain. However, currently, it is unclear to what extent MRS measures reflect stable trait-like neurotransmitter levels, or may be sensitive to the brain’s activity state as well. Therefore, this study investigated if cortical GABA (GABA+/Cr) and Glutamate (Glx/Cr) levels differ as a function of task demand, and if so, in which activity state these measures may best predict behavioral performance. We acquired 3T-MRS data from thirty healthy men in two brain areas during different task demands: the medial occipital cortex (OC), at rest (eyes closed) and while subjects watched a movie (on-task); and the left dorsolateral prefrontal cortex (lDLPFC), at rest, during an easy working memory (WM) task, and during a challenging WM task. Task demand had no effect on the concentration of GABA or Glutamate in either brain region. Moreover, we observed no correlations between GABA and Glutamate concentrations and behavioral performance; occipital neurotransmitter concentrations did not predict visual discrimination nor did those in lDLPFC predict WM updating accuracy, capacity or maintenance. These null findings were supported by Bayesian statistics. In conclusion, these results suggest that with 3T-MRS we measure relatively stable trait-like neurotransmitter concentrations, but at the same time question the validity of 3T-MRS as a method to relate GABA and Glutamate concentrations to behavior.

## Introduction

Magnetic Resonance Spectroscopy (MRS) is a non-ionizing technique that can be used to non-invasively determine in-vivo neurotransmitter concentrations, such as GABA and Glutamate, in the human brain. Being the primary inhibitory and excitatory neurotransmitters, GABA and Glutamate play a key role in regulating neuronal excitability and hence determining cortical functioning. As such, MRS is a promising neuroimaging technique to investigate the relationship between neurotransmitter concentrations and brain functioning and behavior has opened new avenues for investigating the relationship between neurotransmitter concentrations and brain functioning and behavior (Isaacson & Scanziani, 2011).

In recent years, MRS-measured cortical GABA in specific brain areas has been linked to inter-individual differences in related functions in a range of cognitive domains. For example, GABA concentrations in the sensorimotor cortex have shown to be predictive motor performance, as well as tactile discrimination (Puts, Edden, Evans, McGlone, & McGonigle, 2011), while GABA in the occipital cortex (OC) has been related to visual performance (Edden, Muthukumaraswamy, Freeman, & Singh, 2009; Sandberg et al., 2014; Song, Sandberg, Andersen, Blicher, & Rees, 2017; van Loon et al., 2013). Similarly, GABA levels in prefrontal areas have been found to relate to higher-order cognitive functions, such as working memory (Yoon, Grandelis, & Maddock, 2016) and attention (Kihara, Kondo, & Kawahara, 2016). Yet, in many of these studies, sample sizes were relatively small and their findings hence warrant replication.

Moreover, so far, the vast majority of studies linking cortical neurotransmitter levels to behavior have quantified these in rest only, thus assuming that MRS-measured GABA and Glutamate levels reflect stable individual ‘trait’ differences. Yet, other studies that have looked at differences in GABA and Glutamate concentrations as a function of experimental manipulation have found these concentrations may in fact not be so static but can change over relatively short time windows, e.g. as a function of time on task (Michels et al., 2012) or after learning (Floyer-Lea, Wylezinska, Kincses, & Matthews, 2006; Shibata et al., 2017).

At present, it thus remains unclear to what extent MRS-measured neurotransmitter levels reflect stable and consistent, trait-like neurotransmitter concentrations, or in fact are sensitive to changes in metabolic activity as a function of task demand. However, this knowledge is important for our theoretical understanding of what it is that we measure with MRS (‘trait’ or ‘state’ concentrations) and hence has pivotal implications for the design of future studies that aim to experimentally manipulate neurotransmitter concentrations. Moreover, it is still unclear whether neurotransmitter levels that are measured during performance of a task that activates the brain area of interest, are more indicative of performance than neurotransmitter levels measured at rest. Namely, similar to what has been observed with other neuroimaging methods such as EEG and fMRI, neurotransmitter activity investigated in an active state may in fact be a better predictor of behavior than the measures acquired at rest that we currently use in MRS research.

To address these outstanding issues, the current study investigated whether 3T-MRS GABA and Glutamate concentrations vary as a function of task demand, and if so, in which brain state (rest or on-task), these concentrations may best predict cognitive performance. To this end, we scanned both a primary sensory (medial occipital cortex, OC) and higher-order cognitive brain region (left dorsolateral prefrontal cortex lDLPFC).

The OC, key for visual processing was scanned once at rest (eyes closed) and once while subjects watched a movie (on-task). The lDLPFC, which has consistently been shown to be active with temporarily holding and manipulating information in working memory (WM) (Owen, McMillan, Laird, & Bullmore, 2005), was measured three times: at rest, during an easy WM task (letter 2-back), and during a challenging WM task (adaptive letter N-back).

In a separate behavioral session participants performed a visual discrimination task (with oblique grating patches) and two WM tasks (letter N-back updating and Sternberg task). This design critically permitted us to determine, first of all, if MRS-measured GABA and Glutamate levels reflect stable trait-like indicators of brain neurotransmitter concentrations or whether they are influenced by the cognitive state of the subject. Secondly, it allowed us to examine, if concentrations fluctuate with state and task, which state may best predict individual differences in performance outside the scanner.

We expected to replicate previous findings which associated higher occipital GABA levels with better visual discrimination performance (Edden et al., 2009) and higher lateral prefrontal GABA levels with better WM performance (Yoon et al., 2016). We expected no such correlations with Glutamate levels. Moreover, it has recently been proposed that not so much the concentration of each neurotransmitter individually, but the relative concentrations of GABA and Glutamate (i.e. the cortical excitation/inhibition balance) may provide a more accurate reflection of cortical functioning and hence be a better predictor of cognitive performance (Krause & Cohen Kadosh, 2014). In line with this, we expect that combining information from both measures into Glutamate/GABA ratio’s may better predict individual differences in performance than GABA levels only.

## Methods

### Participants

Thirty healthy volunteers (mean age: 21,2 years, SD: 2,5; all men) were recruited via the university subject pool and participated in return for a monetary reward or course credit. As cortical GABA concentrations have shown to vary with the menstrual cycle (De Bondt, De Belder, Vanhevel, Jacquemyn, & Parizel, 2015; Harada, Kubo, Nose, Nishitani, & Matsuda, 2011), only male participants were included. Subjects gave written informed consent before the experiment and the experiment was approved by the University of Amsterdam ethical committee. All reported no history of psychiatric conditions, complied to the rules for MRI safety, and had normal or corrected-to-normal vision.

### Procedure

Subjects came to the lab for two sessions (see Figure 1A), a behavioral and an MRS session, planned at the same time of day with maximally 11 days (mean: 4,2 StD: 3,2) in between. In the first behavioral session, they were seated in a comfortable chair in front of a computer screen (at approximately 90 cm distance) and performed three WM tasks and a visual discrimination task. Order of the tasks was counter-balanced across subjects, and they first practiced each task before data collection started. At the end of the behavioral session, subjects also performed an attentional blink task, but these data were not analyzed for the current paper.

**Figure 1.**
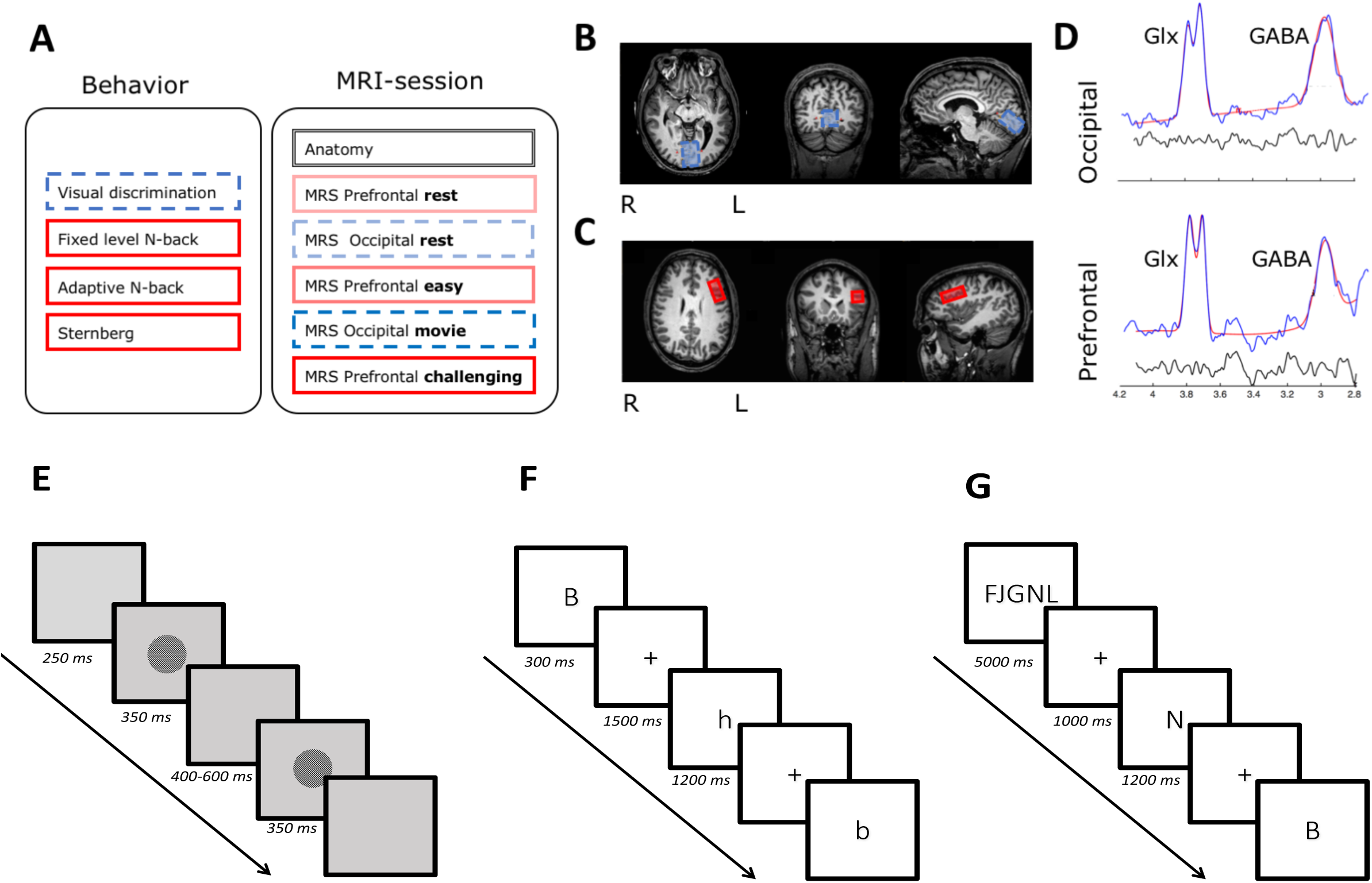
Schematic illustration of the research design and methods. Subjects came to the lab for two sessions, an MRI and behavioral session. (**A**). In the MRI session, 3T-MRS (MEGA-PRESS) was used to measure GABA (GABA+/Cr) and Glutamate (Glx/Cr) levels in an occipital (OC) and a prefrontal voxel (lDLPFC) under different activity conditions. The occipital voxel (**B**) was scanned twice: once when subjects had their eyes closed (rest) and once while they watched a movie (on-task). The prefrontal voxel (**C**) was scanned three times: once with eyes closed (rest), once while subjects performed an easy WM updating task (letter 2-back) and once while they performed a challenging WM updating task (an adaptive letter N-back). Order of the activity conditions was counter-balanced between subjects, but the occipital voxel was always scanned in between the prefrontal voxels. (**D**) Outcome of the modeling of the GABA and Glx signal in the occipital and prefrontal voxel for a typical subject (output from the Gannet analysis toolbox (Edden et al. 2014, www.gabamrs.com). In blue the edited spectrum is shown, overlaid in red is the model of best fit (using a simple gaussian model) and the residual of these is shown in black. (**E-F**): In a separate behavioral session, we administered four tasks to determine cognitive performance. An oblique visual discrimination task (**E**) was performed to relate to neurotransmitter levels in the occipital voxel. Furthermore, three WM tasks (**F**) were administered to relate to neurotransmitter levels in the prefrontal voxel: two versions of the letter N-back WM task (**F**) to determine both WM updating accuracy (level N ranged 2-5) and Capacity (level N on-line adapted to performance) as well as a Sternberg WM maintenance task (**G**).

**Figure 2.**
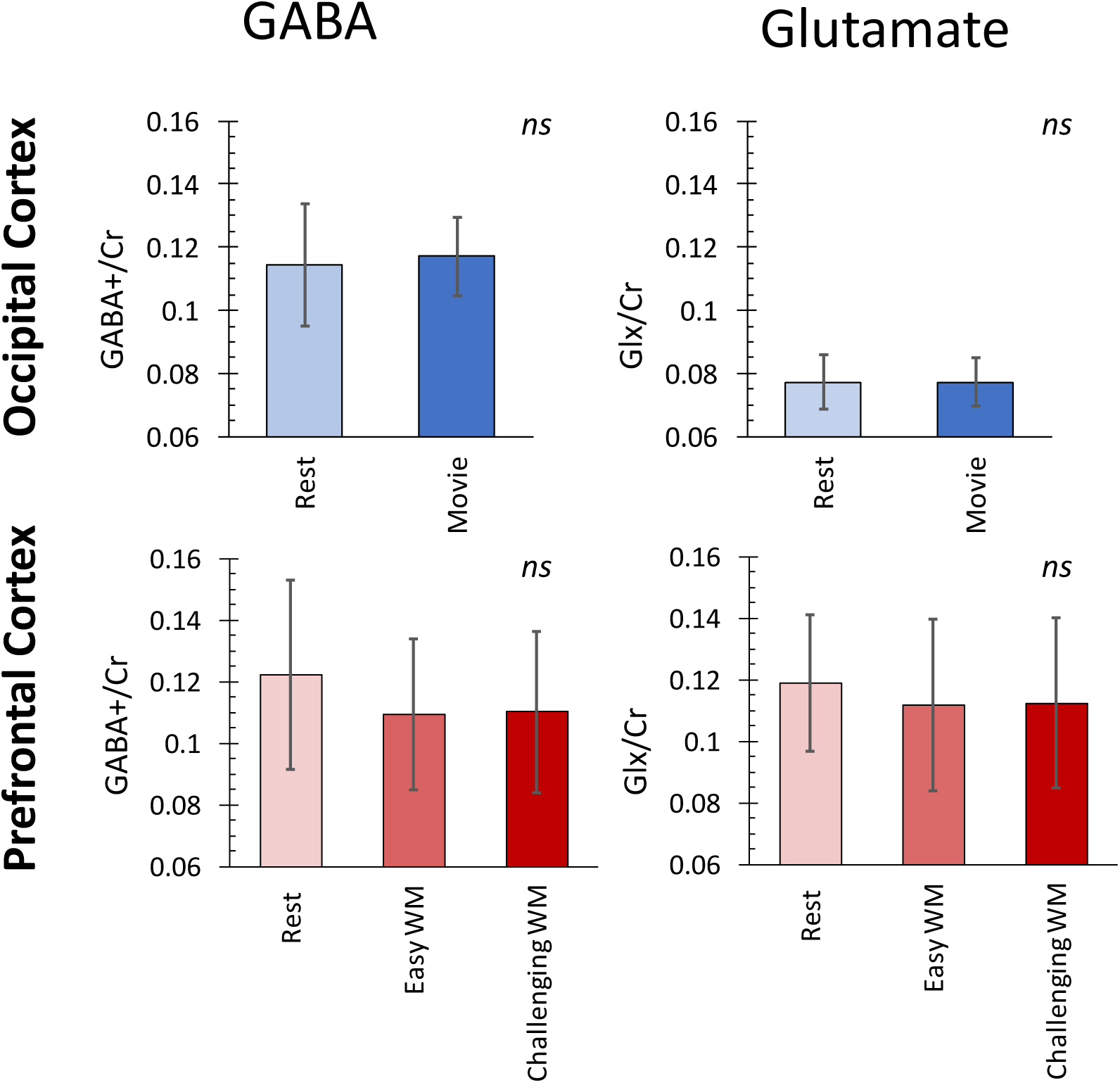
GABA and Glutamate levels did not vary as a function of activity state. Group-level and of GABA (GABA+/Cr) and Glutamate (Glx/Cr) levels per brain area (Occipital cortex: top panel; lDLPFC: bottom panel) and activity condition.

**Figure 3.**
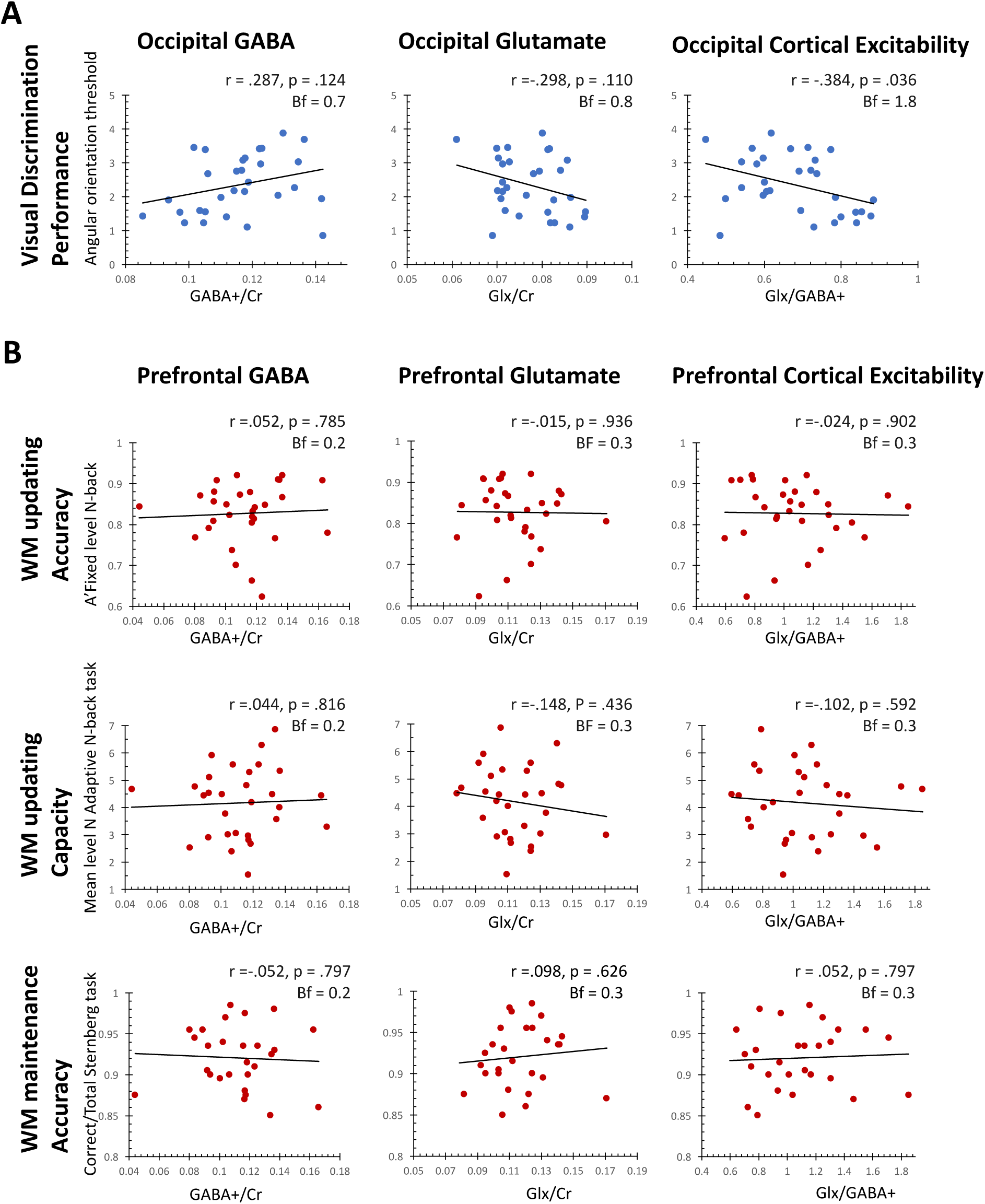
Scatter plots displaying the relationship between GABA and Glutamate concentrations as well as cortical excitability (Glutamate/GABA ratio) (collapsed across activity state) and performance on the brain-region related tasks. None of these metabolite-behavior relationships was significant, indicating that our 3T-MRS measures of GABA and Glutamate did not robustly related to performance outside the scanner. More specifically, occipital cortex neurotransmitter levels, nor cortical excitability predicted visual discrimination performance (**A**), neither did these measures in the lateral prefrontal cortex predict performance on any of the three WM tasks (**B**). Pearson correlation coefficients and two-tailed p statistics are reported (alpha = .025; adjusted for multiple comparisons) as well as Bayes factors.

The visual discrimination task was an orientation task with oblique gratings, similar to the one used by Edden et al. (2009), as described in more detail below. WM performance was measured with two versions of the letter N-back task; one with level N fixed (WM updating accuracy) and one with level N adapted to performance (WM updating capacity). Also, to be able to examine the extent to which metabolite levels in lDLPFC could predict WM performance in a more generalized manner (i.e., on a different WM task than administered during scanning), we furthermore administered a Sternberg task to determine WM maintenance more specifically. Importantly, the Sternberg task has consistently been related to functioning in the lDLPFC in particular, both in functional neuroimaging (Altamura et al., 2007) and non-invasive brain stimulation (Jansma et al., 2013) studies. All three WM tasks are described in more detail below.

In the second MRS session, five MRS scans and an anatomical scan were acquired. In two of the MRS scans, the voxel was placed over the medial occipital cortex (OC) (primary visual cortex, see figure 1B) and in the other three over the left dorsolateral prefrontal cortex lDLPFC (see figure 1C). The OC voxel was scanned twice, once when subjects had their eyes closed (rest condition) and once when they watched a movie (active condition). The lDLPFC voxel was scanned three times: once when subjects had their eyes closed (rest condition), and once while they performed an easy (letter 2-back) and once a challenging (adaptive letter N-back) WM task (see for more info about the task below). By manipulating WM task difficulty for the lDLPFC voxel, we aimed to also investigate possible differences in neurotransmitter levels depending on the extent of cortical engagement. To prevent carry-over effects of task activity in the MRS signal between the different activity states, the lDLPFC and occipital voxels were scanned in an interleaved manner. Also, order of the tasks (and thus activity states) was counter-balanced across subjects. Due to a shortage of time, for one subject, the lDLPFC rest condition scan could not be acquired.

### MRS data acquisition and analysis

Scanning was performed on a 3T Philips Achieva TX MRI scanner (Philips Healthcare) with an eight-channel head coil. Spectroscopy voxel localization was performed by the experimenter according to the individual’s anatomical landmarks as visible from an initial anatomical scan. The occipital voxel (30 x 25 x 20 mm) was placed bilaterally over the calcarine sulcus (see Figure 1B) (cf. van Loon et al., 2013). For the left dorsolateral prefrontal cortex lDLPFC voxel, the center of the voxel (30 × 20 × 25 mm) was placed on the left middle frontal gyrus, with the posterior border of the voxel positioned anterior to the precentral sulcus (see Figure 1C). Both voxels were placed with care to exclude cerebral spinal fluid (CSF) from the ventricles or the cortical surface.

Edited ^1^H J-difference spectra were acquired for each voxel using a GABA-specific sequence of the Mescher-Garwood point-resolved spectroscopy (MEGA-PRESS) method (Waddell, Avison, Joers, & Gore, 2007). Scanning took approximately 12 minutes per acquisition, during which 384 transients were collected (TE = 73 ms; TR = 2,000 ms). On the odd transients, a 15,64 ms sinc-center editing pulse (64 Hz full width at half maximum) was applied in an interleaved manner at 1,9 ppm and 4,6 ppm to excite GABA and suppress water respectively.

Spectral data were analyzed with the MATLAB-based package GANNET v2.1 (Edden et al. 2014, www.gabamrs.com). Using the in-build options of the GannetLoad-function, the following processing steps were performed: time-domain frequency-and-phase correction using spectral correction, line broadening with an exponential apodization function, Fast Fourier Transform (FFT), time averaging, frequency and phase correction based upon fitting of the Choline and Creatine signals, pairwise rejection of the data for which fitting parameters are greater than 3 SDs from the mean, and finally, subtraction of the even from the odd transients to generate the edited difference spectrum. Notably, in this edited difference spectrum, the GABA signal is contaminated by the macromolecule homocarnosine (Edden, Puts, & Barker, 2012), a GABA derivative, and thus often referred to as GABA+. Also, as the spectra of Glutamate and Glutamine are known to overlap at 3T, the combined measure of Glx was used as the best measure for Glutamate.

Subsequently, using the GannetFit function of GANNET, GABA+ and Glx functions were modeled to the data together (see Figure 1D) and ratio’s relative to Creatine (Cr) were calculated (i.e. GABA+/Cr and Glx/Cr). Normalizing values to Creatine has been shown to be superior to normalizing to H2O with regard to intra-subject stability (Bogner et al., 2010; Greenhouse, Noah, Maddock, & Ivry, 2016) and is known to substantially reduce inter-subject variance as a result of differences in global signal strength, as well as those stemming from differences in tissue fractions in the scanned voxel (gray matter, white matter, and cerebrospinal). Calculating GABA and Glutamate levels relative to Creatine thus makes coregistration, segmentation and the calculation of CSF corrected values superfluous. Scans were excluded when no Creatine peak was visible in the data (N=3; corresponding to Creatine Signal to Noise ration (SNR) < 50), model fit turned out to be poor (N=2; GABAGlxModelfiterror > 15), or the GABA+ or Glx peak could not be confidently be determined (N=1; GABA SNR < 3). Furthermore, we used the Statistical Package for the Social Sciences for Mac OS, Version 24 (IBM, Armonk, NY) to identify outliers as a result two GABA+/Cr values in the lDLPFC rest condition (0.359, 0.238) and three in the easy WM condition (0.347, 0.246, 0.201) as extreme outliers. As these outlier values were also much higher than previously reported in the prefrontal cortex (De Bondt et al., 2015; Greenhouse et al., 2016), these were excluded from further analyses. The remaining GABA+/Cr and Glx/Cr ratios in the OC and lDLPFC voxels fell all in agreement with previous studies that measured similar regions during rest (De Bondt et al., 2015; Edden et al., 2009; Greenhouse et al., 2016; Iwabuchi et al., 2017; Michels et al., 2012; Yoon et al., 2016).

### Visual discrimination task

In the behavioral session only, participants performed a visual discrimination task that was based on the one used by Edden and colleagues (2009). In this task, subjects were sequentially shown two circles with oblique grating patterns and asked to indicate if the second of the two was rotated clockwise (left mouse button) or counterclockwise (right mouse button) with respect to the first one (see Figure 1E). The circular gratings (diameter: 4 degrees visual angle, Spatial frequency: 3 cycles/degree, contrast: 80%, mean luminance: 44,5 cd/m2) were displayed for 350 ms each, with an inter stimulus interval chosen randomly between 400 and 600 ms. During the task, the difference in orientation was adjusted logarithmically, using two interleaved staircases that applied the principle of one up, two down. Mean orientation of both gratings was always 45 degrees, as Edden et al. (2009) observed the highest correlation between GABA and orientation discrimination threshold in an oblique compared to a vertical average condition. An auditory tone provided feedback on each trial, and one run of the task continued until both staircases completed 12 reversals. Subjects completed two runs of the task, but only the second run was used for analysis due to expected early task training effects (as reported by Edden et al., 2009). Of this second run, the first two reversals were discarded and visual discrimination thresholds were subsequently computed for each participant by averaging the angle difference between the two stimuli over the last 10 reversals and both staircases (cf. Edden et al., 2009).

### Working memory tasks

The primary working memory (WM) task that subjects performed in both the behavioral and MRS session was a letter N-back task (see Figure 1F). In this WM updating task, subjects are presented with a stream of letters and asked to indicate if the currently presented letter is the same as the one presented N stimuli back. Hereby, N is an integer and the value of N determines the difficulty level of the task. With higher levels of N, more stimuli have to be held in WM in sequential order, increasing WM load. As WM content has to be continuously updated, the letter N-back task is considered to be a demanding WM task. Therefore it is a standard task to investigate WM updating performance (Jaeggi, Buschkuehl, Perrig, & Meier, 2010) and importantly, has consistently been related to processing in the lDLPFC (e.g., see meta-analysis by Owen et al., 2005).

In our letter N-back task letters (Arial, font size 72, letterset [“A”, “B”, “C”, “D”, “E”, “F”, “G”, “H”, “J”, “K”]) were presented for 300 ms at the center of a screen, followed by a 1500 ms inter-stimulus interval in which a fixation cross was displayed (Arial, font size 20). In the behavioral session, we presented black letters on a white screen, while in the MRS session we showed subjects white letters on a black background because of the dimly lit nature of the scanning room. Of the presented letters, approximately 35% were so-called targets, i.e., the letter in the current trial matched the letter presented N letters back. Letters could be presented in upper or lower case and would still classify as the same letter (i.e., a target). If presented with a target, subjects were required to press the space bar on the keyboard in front of them in the behavioral session, or one of the buttons on the button-box in the scanner. Runs consisted of a stream of 20 + N stimuli each and were self-paced in the behavioral session to allow the subject to take small breaks in between and enhance focus during the runs, but they started automatically in the MRI-session to ensure the task was performed for the entire time of the scan.

Subjects performed both a fixed-level and an adaptive version of the letter N-back task in the behavioral and in the MRS session. First of all, in the behavioral session, subjects performed 24 runs of a fixed-level version of the task in which level N sequentially increased from 2 to 5, with steps of 1. We used this version of the task to calculate WM updating accuracy, which was operationalized using A’ (A prime). A’ is the non-parametric variant of signal detection theory’s d’ and takes into account both hits (correct responses) and false alarms (incorrect responses). In contrast to d’, A’ can account for situations in which participants do not show any false alarms, which sometimes occurred on lower N levels of our task. A’ scores range from 0 to 1, with 0 indicating chance performance and 1 perfect accuracy. A’ can be calculated from hit rate (H) and false alarm rate (F) with the following formula (Zhang & Mueller, 2005):

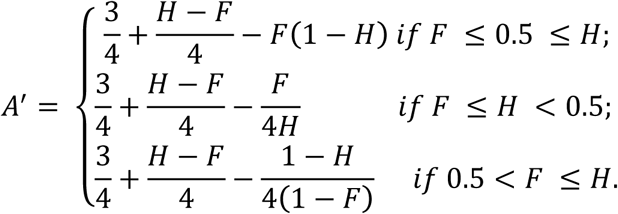

Secondly, in the adaptive version of the task, level of N always started with N = 2 (set as the lowest possible level N) and subsequently adapted to performance by going up one step (current N+1) if subjects made fewer than three errors, and down one step (current N-1) if they may more than five errors (similar to Jaeggi, Buschkuehl, Jonides, & Perrig, 2008). The adaptive version of the task in the behavioral session also consisted of 24 runs, of which we calculated average level N over the last 21 runs only (disregarding the first 3 runs to allow each individual some ramp-up time to their average level), to use as our measure for WM updating capacity. Additionally, in the MRS-session, subjects once performed the task with N fixed to level 2 (the *easy* WM condition) and once with N adapted to performance in the same way as in the behavioral session (the *challenging* WM condition). Hereby, the amount of runs of the task was determined to ensure it covered the whole MRS-scan. We used Presentation software (Neurobehavioral Systems, Inc.) to administer the letter N-back task.

In the Sternberg task (see Figure 1G), subjects were presented with a string of five letters that they were required to remember (5000 ms). Consequently, one letter was shown on the screen at a time (1200 ms per letter, 1000 ms fixation cross in between) for which subjects had to indicate whether that letter was in the currently remembered string (press ‘N’ key) or not (press ‘Z’ key). All letters were presented in uppercase and came from a predetermined letterset ([“B”, “D”, “F”, “G”, “H”, “J”, “K”, “L”, “M”, “N”], Arial, Fontsize 60). Per run, 10 letters were presented of which 50% was a target. Subjects completed 10 runs of the task. We determined WM maintenance accuracy by calculating the number of correct trials divided over the total. The data of one subject was discarded because of extreme low below chance performance (accuracy = 30% correct).

### Specific hypotheses and statistical approach

To investigate whether GABA and Glutamate levels measured with MRS differed between different levels of task demand, repeated measures ANOVA’s were run with task demand as the within-subjects factor and GABA+/Cr or Glx/Cr as the dependent variable for the occipital and lDLPFC voxel separately. Additionally, to investigate within-subject stability of MRS measured GABA and Glutamate levels across activity states, after testing for normality, individual Pearson correlations were run between the GABA+/Cr and Glx/Cr levels measured under the different task conditions, again separately for the prefrontal and occipital brain region. Namely, we reasoned that if these neurotransmitter measures reflect stable, trait-like neurotransmitter concentrations, they should correlate across activity states across subjects. Furthermore, we wanted to replicate previous reports of within-subject regional specificity for GABA-levels (Bogner et al., 2010; Greenhouse et al., 2016), therefore we also correlated occipital and lDLPFC neurotransmitter measures in the resting state.

Depending on the outcomes of our first ANOVA’s, we followed one of two approaches. In case of no specific effect of activity state, GABA+/Cr and Glx/Cr concentrations were averaged over all conditions separately for the OC and lDLPFC and these average measures were related to individual task performance; OC: rest and movie, DLPFC: rest, easy (letter 2-back) and challenging (adaptive N-back). In case of systematic differences in MRS measures between activity states, multiple regression analyses including all activity states as predictors were run to determine which activity state best predicted behavioral performance. Analyses were conducted separately for GABA and Glutamate and included the relevant brain area and corresponding task only. In addition, we also ran control analyses in which neurotransmitter concentrations of the task-unrelated brain region were related to the behavioral measures, for which we did not expect to find significant correlations.

To test the hypothesis that not so much GABA and Glutamate individually, but in fact their relative concentrations (i.e. the excitation/inhibition balance) may provide the best predictor of cognitive functioning, for both brain areas we also calculated glutamate/GABA ratio’s from our data and investigated the extent to which these ratio’s predicted behavioral performance.

The correlation with our behavioral measures were investigated with 2-tailed Pearson correlations. To account for the fact that GABA and the Glutamate/GABA ratio are highly related and investigate a similar research question, we divided alpha over two to determine significant levels and correct for multiple comparisons.

All statistical analyses were conducted using the Statistical Package for the Social Sciences for Mac OS, Version 24 (IBM, Armonk, NY). Furthermore, we additionally repeated our analyses with Bayesian statistics using the open-software package JASP (http://www.jasp-stats.org, Wagenmakers, Marsman, et al., 2017)). The resulting Bayes factors, which grade the intensity of evidence for the null (H0) and alternative hypothesis (H1), and values were interpreted according to the corresponding classification scheme (see for elaboration Wagenmakers, Love, et al., 2017): 1/30 < Bf < 1/10, Strong evidence for H0; 1/10 < Bf < 1/3, Moderate evidence for H0; 1/3 < Bf < 1, Anecdotal evidence for H0; Bf = 1, No evidence; 1 < Bf < 3, Anecdotal evidence for H1; 3 < Bf < 10, Moderate evidence for H1; 10 < Bf < 30, Strong evidence for H1.

## Results

### Descriptives cognitive performance

Subjects performed in line with expectations on all tasks in both the behavioral and the MRS session. Visual discrimination angle thresholds ranged between 0.845 and 3.873 (Mean: 2.347, StD: 0.868), similar to the range reported by e.g., Edden et al. (2009). For the WM tasks, accuracy was well above chance for all participants on both the fixed WM updating letter N-back task (range A’: 0.623 to 0.920, mean: 0.827, StD: 0.075), and the Sternberg maintenance task (range Accuracy: 0.850 to 0.950, mean: 0.921, StD: 0.038). Moreover, on the adaptive N-back task, subjects showed a relatively wide spread in WM updating capacity (range mean level N: 1.53 to 6.86, mean: 4.17, StD: 1.28), i.e., inter-individual differences in WM updating capacity were relatively large.

In the MRS session, accuracy on the 2-letter N-back task ranged between 0.79 and 1.00 (A’ Mean: 0.95, StD: 0.04), indicating ceiling level or close to ceiling level performance in all subjects. Also, WM capacity levels on the adaptive N-back task were similar to those observed in the behavioral session (range mean level N: 2.20 to 7.80, Mean: 4.54, StD: 1.50) and correlated well within subjects (r = .826, p < .001; Bf = 743996). These findings show that our task manipulation was effective, with the adaptive WM task (i.e. challenging WM condition) placing greater demands on WM processes than the 2-letter N-back task (i.e. easy WM condition).

### Resting-state versus on-task occipital and prefrontal GABA and Glutamate levels

To address our first research question, we assessed whether MRS-measured GABA and Glutamate levels differed as a function of task demand (i.e. reflect activity state). To this end, we compared neurotransmitter levels measured in rest with those measured during stimulus-or task-induced activity, separately for the OC and lDLPFC voxels.

In the occipital voxel, we observed no difference in GABA or Glutamate concentrations between the rest and active (movie watching) condition (GABA: (F(1,29) = .904, p = .350, Bf = 0.37) and Glutamate: (F(1,29) < .001 p = .981, Bf = 0.26). Similarly, GABA levels in lDLPFC did not show significant differences between the three activity conditions (Rest; Easy WM and Challenging WM) (F(2,38) = .210, p = .811, Bf = 0.16) and neither did Glutamate (F(2,38) = .210, p = .811, Bf = 0.30). Thus, both in the occipital and in the prefrontal cortex we found no effect of task demand on GABA and Glutamate concentrations. Importantly, in all these cases our Bayesian statistics showed moderate evidence for the null-hypothesis. Together, these findings thus indicate that our MRS measure was insensitive to possible stimulus-or task-induced changes in GABA and Glutamate levels.

Additionally, within subjects GABA levels correlated well between the Rest and Movie conditions in OC (r(29)= .568 p < .001, Bf = 37.8) as well as between all task demand conditions in lDLPFC (Rest and Easy WM: r(21) = .400 p = .032, Bf = 1.3, Rest and Challenging WM: r(25) = .544, p = .002, Bf = 10.4, Easy and Challenging: r(20) = .410, p = .032, Bf = 1.4). Our Bayesian statistics thus indicate strong overall intra-subject stability of GABA levels in the OC, but only anecdotal evidence for intra-subject stability of GABA levels in the lDLPFC. Similarly, Glutamate levels significantly correlated between the two conditions in OC (r(29) = .531, p = .003, Bf = 17.5. However, in lDLPFC, Glutamate correlated well between the Rest and Easy WM (r(25) = .476, p = .014, Bf = 4.287), but not between the Rest and the Challenging WM (r(26) = .157, p = .434, Bf = 0.3) and the easy and the Challenging WM (r(23) = -.120, p = .575, Bf = 0.3) conditions. In this case, our Bayesian statistics produce a similar picture, providing strong evidence for the within-region intra-subject stability of our Glutamate measure in the occipital cortex and anecdotal to moderate evidence for within-region intra-subject stability of our Glutamate measure in lDLPFC.

Replicating previous findings of regional specificity of neurotransmitter levels (Bogner et al., 2010; Greenhouse et al., 2016), resting-state GABA levels did not correlate between the OC and lDLPFC voxel (r(26) = -.256, p = .198, Bf = 0.5), nor did Glutamate levels (r(28) = .124, p = .521, Bf = 0.3).

In sum, GABA and Glutamate levels did not systematically change depending on the activity state of the brain region but were relatively stable over the different task demand conditions within subjects. This indicates that although current 3T-MRS neurotransmitter concentrations do not capture possible differences in neurotransmitter activity between activity states (rest versus on-task), they do reliably capture stable trait-like measures of individual neurotransmitter levels in the human brain, especially with regard to GABA and in the occipital cortex.

### Linking occipital and prefrontal GABA and Glutamate to region-related cognitive performance

Our second main aim was to determine, if MRS-measured neurotransmitter concentrations were sensitive to the activity state of the brain region, in which activity state the concentrations would best predict individual differences in behavioral performance. However, given that there were no significant differences between activity states (on task, rest), we continued by averaging GABA and Glutamate per brain region over the different task demand conditions.

#### Occipital GABA and Glutamate and visual discrimination performance

First, we related occipital GABA and Glutamate levels to visual discrimination performance. We expected to replicate the negative correlation between resting-state occipital GABA levels and visual discrimination performance previously reported by Edden et al. (2009), but no correlation between Glutamate and visual discrimination performance. However, in contrast to our expectations and the findings by Edden et al. (2009), participants’ average GABA levels in OC did not predict their performance on the visual discrimination task (r(29) = .287, p = .124, Bf = 0.7). In line with our expectations, average Glutamate levels in OC did not either (r(29) =-.298, p = .110; Bf = 0.8). In both cases, Bayesian statistics reported anecdotal support for the null hypothesis of no relationship. When using the Glutamate/GABA ratio’s as an index of cortical excitability (Krause, Márquez-Ruiz, & Kadosh, 2013), we observed a correlation, namely higher Glutamate/GABA ratio’s correlated with lower discrimination thresholds (r(29) = -.384, p = .036; Bf = 1.8), but this correlation does not survive our multiple comparison correction and is backed up with only anecdotal evidence according to Bayesian statistics.

A post-hoc analysis revealed that even when we correlated GABA levels only at rest like Edden et al. (2009) (i.e., not averaged across conditions (Rest and Movie)), we observed a trend-level correlation also in the opposite direction (r(29) = .328, p = .077, Bf = 1.0). However, this correlation would again not survive a multiple comparison correction and moreover was supported with zero to no evidence according to our Bayesian results.

Thus, contrary to our expectations, we conclude that occipital cortex GABA, Glutamate and cortical excitability levels were not related to visual discrimination performance in our study.

#### Prefrontal GABA and Glutamate and WM performance

Next, we examined the relationship between GABA and Glutamate levels in the lDLPFC and behavioral performance on the three WM tasks performed in the separate behavioral session. In line with Yoon et al. (2016), we predicted that higher lDLPFC GABA levels would predict better WM performance. As Yoon et al. (2016) specifically found a correlation between prefrontal resting-state GABA and performance degradation as a result of increased WM load, but not increased maintenance time or as a function of distractor presence, we furthermore expected that this relation would be specifically apparent for our WM updating capacity measure, as this measure may be the most sensitive to inter-individual differences in WM load. We did not expect lDLPFC Glutamate levels to significantly predict WM performance, nor did we expect occipital GABA levels to predict WM performance (both also similar to Yoon et al., 2016).

In contrast to expectations, but mirroring the OC results, average GABA levels in the lDLPFC did not predict accuracy on the fixed level Letter N-back task (r(29) = .052, p = .785, Bf = 0.2), mean level N on the adapted N-back (r(29) = .044, p = .816, Bf = 0.2), or accuracy on the Sternberg maintenance task (r(27) = -.052, p = .797, Bf = 0.2). In line with our expectations, lDLPFC Glutamate levels did not either (WM updating accuracy: r(29) = -.015, p = .936, Bf = 0.3; WM capacity: r(29) = -.148, p = .436, Bf = 0.3; WM maintenance (Sternberg): r(27) = .098, p = .626, Bf = 0.3). Furthermore, we looked at the Glutamate/GABA ratio as a possibly more sensitive index of cortical excitability, but this measure also did not significantly correlate with WM updating accuracy (r(29) = -.024, p = .902, Bf = 0.3), WM capacity (r(29) = -.102, p = .592, Bf = 0.3), nor WM maintenance (r(27) = .052, p = .797, Bf = 0.3). In all cases, Bayesian analyses indicated moderate evidence for the null hypothesis that the lDLPFC neurotransmitter measures do not relate WM performance. Even when we reran all analyses, but looked at resting-state only to stay closest to current literature (Yoon et al., 2016), no significant relationship between neurotransmitter levels or cortical excitability and WM performance was observed for any of our three WM tasks (all p’s > .139, all Bf’s < 0.7).

We reasoned that perhaps the delay between the behavioral session and the MRS session could have decreased our sensitivity to neurotransmitter concentration and brain-behavior correlations. Therefore, post-hoc, we also explored these correlations with WM performance measured during the MRS scanning procedure. This, however, produced qualitatively the same pattern of null findings: nor GABA, nor Glutamate, nor cortical excitability measured during the Easy WM task could predict simultaneously acquired accuracy scores on this 2-letter N-back task (all p’s > .142, Bf’s < 0.7). Similarly, neurotransmitter levels acquired during the Challenging WM task did not predict WM updating capacity as measured during scanning either (all p’s > .279, Bf < 0.5).

In summary, in contrast to our predictions, the neurotransmitter concentrations we measured in the occipital cortex and the left dorsolateral prefrontal cortex did not correlate with visual discrimination and WM performance, respectively. Thus, while our first set of findings suggested that GABA and Glutamate levels measured with 3T-MRS may reflect relatively stable measures of individual neurotransmitter concentrations, they seem to fail to predict individual differences in behavioral performance on brain region-relevant tasks. Remarkably, we thus did not replicate previous reports of such relationships, even though our sample size was substantially larger than in both of these studies (30 versus 15 and 23 respectively (Edden et al., 2009; Yoon et al., 2016)).

## Discussion

The current study set out to investigate to what extent 3T-MRS measured GABA and Glutamate levels capture changes in cognitive activity state, as well as to determine under which activity state (rest vs. on task) these concentrations may best predict behavioral performance. We observed no differences in GABA or Glutamate levels during resting state compared to active, on-task conditions, neither in the primary visual cortex (the occipital cortex) nor in a higher-order prefrontal area (left DLPFC). Importantly, in general, we did observe strong within-subject correlations between the GABA and Glutamate levels for the different conditions within each brain area, showing that the measurements themselves where reliable. Furthermore, in contrast to previous findings, in this study levels of GABA and Glutamate, or their ratio (averaged over activity states), did not predict inter-individual differences in behavioral performance on brain region-related cognitive tasks.

Together, these findings therefore suggest that 3T-MRS may provide relatively stable ‘trait’-like measures of GABA and Glutamate at the neurochemical level which are insensitive to subtle functionally-related changes as a function of cortical activation. At the same time, however, they question a robust relation between these trait-like neurotransmitter concentrations and behavioral individual differences in brain region-related cognitive performance.

Our finding that current 3T-MRS measures of GABA and Glutamate are insensitive to task demand and reflect stable ‘trait’ rather than ‘state’ levels has important implications for the interpretation of previous studies that did observe changes in GABA over relatively short time-windows. More specifically, these studies have consistently reported decreases in GABA concentrations over time; in the sensorimotor cortex after thirty minutes of performance on a motor task (Floyer-Lea et al., 2006), in the occipital cortex after twenty minutes of performance on a visual perceptual learning task (Shibata et al., 2017), and in prefrontal regions after forty minutes of performance on working memory task (Michels et al., 2012). The fact that we did not observe any activity-related changes in GABA in two of these three brain regions suggests that these earlier findings cannot simply be explained by transient modulations in activation because of longer time spent on the task and thus likely reflect learning-related structural changes in GABA activity. Also, indirectly, this implies that 3T-MRS may be a useful method to investigate the role of GABA in such learning-related cortical plasticity, as these changes seem substantial enough to be picked up with this measure.

In both the occipital and prefrontal brain region, GABA and Glutamate levels correlated strongly within subjects between the rest and task activity conditions (except for prefrontal Glutamate in the Challenging WM condition). This indicates that our measures are reliable and relatively stable within subjects. Yet, the obtained correlation coefficients are somewhat lower than previously reported in studies that looked at resting state blocks only (e.g. (Bogner et al., 2010; Greenhouse et al., 2016)). This could suggest that very subtle differential effects of cognitive activity on GABA and Glutamate across individuals may be picked up by our measure. In line with this, a recent 7T-MRS study did not observe any changes in GABA or glutamate as a function of acute psychosocial stress (Hoetepen et al., 2017). In this study, GABA and Glutamate levels were significantly correlated over time in the control condition, but were not correlated in the stress condition. These observations support the notion that activity state (in this case, stress) could indeed have very small, differential effects on GABA and glutamate across individuals. Because of these inter-individual differences, in current MRS practices, these ‘state’-related fluctuations may fail to become visible at the group-level.

Although the measured GABA and Glutamate levels were found to reflect stable, ‘trait’-like neurotransmitter concentrations, we observed no relationship between these levels and individual differences in behavioral performance on region-related tasks. More specifically, in contrast to a previous study by Edden et al. (2009), in our study occipital GABA (both when averaged over conditions and when only looked at rest) did not predict visual discrimination performance. This was unexpected, as we used the same task, observed a similar spread in subject’s performance, and included an MRS voxel that covered a highly similar area of the visual cortex as Edden et al. Considering the relatively small sample size of the previous study (N=15) and only moderate sample size of the current study (N=30), future replication studies with larger sample sizes and thus greater statistical power may be necessary to further investigate the possible absence or presence of a relation between occipital GABA and visual discrimination performance. Indeed, our Bayesian correlation analyses suggested that even with our relatively large sample size, evidence for the null hypothesis of no relationship was only anecdotal.

Mirroring the occipital cortex findings, lateral dorsolateral prefrontal GABA did not predict performance on any of the three WM tasks; measuring WM updating, accuracy, WM capacity as well as WM maintenance. In this case, our Bayesian correlation analyses suggested moderate evidence in our data for the absence of such relationships. Here too, we thus failed to replicate findings by a previous study (Yoon et al. 2016, N=23) in which resting-state lateral prefrontal GABA levels correlated with individual differences on a face WM maintenance task. More specifically, in this study, Yoon et al. found that prefrontal GABA correlated positively with the extent to which subjects’ performance decreased as WM load increased (one versus two to be remembered faces) Yet, no correlations with GABA were found when Yoon and colleagues looked at WM performance differences as a result of increased maintenance time or the absence or presence of distractors, indicating that this relation may hold only for a rather specific aspect of WM. Although the three WM tasks included in the current study are different than the tasks used by Yoon et al., both WM updating and maintenance have been robustly associated with activation of the lDLPFC and may thus be considered region-related WM functions (Altamura et al., 2007; Owen et al., 2005). As the current study included a larger subject sample (N=30), and applied a more extended range of WM tasks, our null results at the very least suggest that the previously reported relationship between WM performance and GABA concentrations in lDLPFC is not very robust. Furthermore, together with the lack of a neurotransmitter-behavior relation in the occipital cortex, they cast doubt on the claim that with current 3T-MRS practices we can detect relationships between neurotransmitters levels and region-related behavioral performance.

One important direction for future studies may therefore be to examine the role of neurotransmitters in cognitive functions using 7T-instead of 3T-MRS. Although less widely available, 7T has two important advantages over 3T with regard to MRS. Firstly, increased spectral resolution at the 7T-MRS enables better discrimination and quantification of neurotransmitter concentrations of both Glutamate (independent from Glutamine (An et al. 2014)) and GABA (uncontaminated by macromolecules (Ganji et al. 2014)). Secondly, at higher field strengths, better signal-to-noise ratio’s may be obtained (Choi et al., 2010), which enables the use of smaller sized MRS voxels, thereby increasing sensitivity to study a precise target region.

Namely, an important limitation of current 3T-MRS is the relatively large MRS voxelsize that is necessary to acquire sufficient signal strength. Placing this relatively large voxel common over an actually much smaller region of interest may substantially ‘delute’ the signal, as small differences in the relevant cortical region (i.e. the desired signal) may drown in a sea of irrelevant fluctuations in the surrounding cortical regions (i.e. noise) that are also included in the voxel and thus together create the average that we measure. In other words, measuring GABA and Glutamate concentrations in a voxel that is much larger than the relevant brain region may substantially reduce the sensitivity of the method to investigate small-scale relevant regional specific neurotransmitter concentrations to relate to behavior. Future studies should therefore investigate if the higher spectral and spatial resolution of 7T-MRS may create a method that is more sensitive to detect changes in neurotransmitter activity induced by task demand, as well as investigate relationships between neurotransmitter function in a specific brain region and related cognitive and behavioral performance.

Another direction which may aid in increasing sensitivity of MRS to detect local neurotransmitter concentrations may be to combine MRS with functional neuroimaging. More specifically, localizing individual peak activations for the region of interest may significantly help to increase spatial acuity in the placement of the MRS voxel over the relevant area. This may be particularly helpful for higher order cortical areas, including the prefrontal cortex, where variability in functional neuroanatomy is particularly high. For example, peak activations on a WM maintenance task are known to be spread along the middle frontal gyrus across individual subjects (Jansma et al., 2013), and thus, a one-fits-all approach here may be less effective. The (relatively large) voxel used in the current study ensures peak activation was covered for all subjects, but conceivably also included surrounding cortical regions not engaged by our tasks. Eventually, therefore, smaller voxels such as may be enabled by higher magnetic field strengths, that are placed individually after functional localization, may enhance spatial acuity substantially and thereby result in higher sensitivity and more accurate measures of neurotransmitter concentrations for a specific functional region of interest.

A last explanation for the lack of brain-behavior correlations in the current study is that the hypothesized relation between neurotransmitter levels and functional performance is actually more complex than the standard simple linear correlation we generally apply to investigate such inter-individual correlations. In fact, with regard to the excitation/inhibition balance, it has been proposed that an inverted U-curve may best describe the relation with performance, with performance being highest when the cortex is active enough for functional firing to effectively take place, but at the same time inhibited enough to reduce noise and unwanted firing (Krause & Cohen Kadosh, 2014). However, to adequately investigate this, many more data points are needed than are currently generally available in neuroimaging studies. This, again, calls for the use of larger sample sizes in studies that attempt to link neurotransmitter levels to behavior.

## Conclusion

To conclude, the current study found that 3T-MRS measures of GABA and Glutamate generally reflect stable and reliable ‘trait’-like neurotransmitter levels and do not capture task demand-induced changes. However, in contrast to previous findings, we did not observe correlations of neurotransmitter concentrations with behavioral performance on region-related tasks. This questions to what extent GABA and Glutamate concentrations measured with current 3T-MRS practices reflect neurotransmitter activity that is relevant for behavior. The use of higher magnetic field strengths (e.g., 7T), and/or individually localized voxel placement in future studies may improve the sensitivity to subtle task-induced changes in GABA and Glutamate levels, allowing further investigation of in-vivo measured neurotransmitter levels in the human brain as well as their relationship with behavior.

## Acknowledgements

We thank Martijn Jansma for sharing his functional activation findings with the Sternberg task with us and Nick Puts for his expertise help with analyzing our data with the GANNET toolbox and his personal efforts to investigate the data quality and Prof. Dr. K. Richard Ridderinkhof for his helpful comments on an earlier version of the manuscript.

## References

Altamura et al. (2007). Dissociating the effects of Sternberg working memory demands in prefrontal cortex. Psychiatry Research: Neuroimaging, 154(2), 103–114. https://doi.org/10.1016/j.pscychresns.2006.08.002

Bogner, W., Gruber, S., Doelken, M., Stadlbauer, A., Ganslandt, O., Boettcher, U., … Hammen, T. (2010). In vivo quantification of intracerebral GABA by single-voxel 1H-MRS—How reproducible are the results? European Journal of Radiology, 73(3), 526–531. https://doi.org/10.1016/j.ejrad.2009.01.014

Choi et al. (n.d.). Improvement of resolution for brain coupled metabolites by optimized 1H MRS at 7 T [Wiley Online Library]. Retrieved November 29, 2017, from http://onlinelibrary.wiley.com/doi/10.1002/nbm.1529/full

De Bondt, T., De Belder, F., Vanhevel, F., Jacquemyn, Y., & Parizel, P. M. (2015). Prefrontal GABA concentration changes in women—Influence of menstrual cycle phase, hormonal contraceptive use, and correlation with premenstrual symptoms. Brain Research, 1597, 129–138. https://doi.org/10.1016/j.brainres.2014.11.051

Edden, R. A. E., Muthukumaraswamy, S. D., Freeman, T. C. A., & Singh, K. D. (2009). Orientation Discrimination Performance Is Predicted by GABA Concentration and Gamma Oscillation Frequency in Human Primary Visual Cortex. The Journal of Neuroscience, 29(50), 15721–15726. https://doi.org/10.1523/JNEUROSCI.4426-09.2009

Edden, R. A. E., Puts, N. A. J., & Barker, P. B. (2012). Macromolecule-suppressed GABA-edited magnetic resonance spectroscopy at 3T. Magnetic Resonance in Medicine, 68(3), 657–661. https://doi.org/10.1002/mrm.24391

Floyer-Lea, A., Wylezinska, M., Kincses, T., & Matthews, P. M. (2006). Rapid Modulation of GABA Concentration in Human Sensorimotor Cortex During Motor Learning. Journal of Neurophysiology, 95(3), 1639–1644. https://doi.org/10.1152/jn.00346.2005

Greenhouse, I., Noah, S., Maddock, R. J., & Ivry, R. B. (2016). Individual differences in GABA content are reliable but are not uniform across the human cortex. NeuroImage, 139, 1–7. https://doi.org/10.1016/j.neuroimage.2016.06.007

Harada, M., Kubo, H., Nose, A., Nishitani, H., & Matsuda, T. (2011). Measurement of variation in the human cerebral GABA level by in vivo MEGA-editing proton MR spectroscopy using a clinical 3 T instrument and its dependence on brain region and the female menstrual cycle. Human Brain Mapping, 32(5), 828–833. https://doi.org/10.1002/hbm.21086

Isaacson, J. S., & Scanziani, M. (2011). How Inhibition Shapes Cortical Activity. Neuron, 72(2), 231–243. https://doi.org/10.1016/j.neuron.2011.09.027

Iwabuchi, S. J., Raschke, F., Auer, D. P., Liddle, P. F., Lankappa, S. T., & Palaniyappan, L. (2017). Targeted transcranial theta-burst stimulation alters fronto-insular network and prefrontal GABA. NeuroImage, 146, 395–403. https://doi.org/10.1016/j.neuroimage.2016.09.043

Jaeggi, S. M., Buschkuehl, M., Jonides, J., & Perrig, W. J. (2008). Improving fluid intelligence with training on working memory. Proceedings of the National Academy of Sciences. https://doi.org/10.1073/pnas.0801268105

Jaeggi, S. M., Buschkuehl, M., Perrig, W. J., & Meier, B. (2010). The concurrent validity of the N-back task as a working memory measure. Memory, 18(4), 394–412. https://doi.org/10.1080/09658211003702171

Jansma, J. M., van Raalten, T. R., Boessen, R., Neggers, S. F. W., Jacobs, R. H. A. H., Kahn, R. S., & Ramsey, N. F. (2013). fMRI Guided rTMS Evidence for Reduced Left Prefrontal Involvement after Task Practice. PLoS ONE, 8(12). https://doi.org/10.1371/journal.pone.0080256

Kihara, K., Kondo, H. M., & Kawahara, J. I. (2016). Differential Contributions of GABA Concentration in Frontal and Parietal Regions to Individual Differences in Attentional Blink. Journal of Neuroscience, 36(34), 8895–8901. https://doi.org/10.1523/JNEUROSCI.0764-16.2016

Krause, B., & Cohen Kadosh, R. (2014). Not all brains are created equal: the relevance of individual differences in responsiveness to transcranial electrical stimulation. Frontiers in Systems Neuroscience, 8. https://doi.org/10.3389/fnsys.2014.00025

Krause, B., Márquez-Ruiz, J., & Kadosh, R. C. (2013). The effect of transcranial direct current stimulation: a role for cortical excitation/inhibition balance? Frontiers in Human Neuroscience, 7. https://doi.org/10.3389/fnhum.2013.00602

Michels, L., Martin, E., Klaver, P., Edden, R., Zelaya, F., Lythgoe, D. J., … O’Gorman, R. L. (2012). Frontal GABA Levels Change during Working Memory. PLOS ONE, 7(4), e31933. https://doi.org/10.1371/journal.pone.0031933

Owen, A. M., McMillan, K. M., Laird, A. R., & Bullmore, E. (2005). N-back working memory paradigm: A meta-analysis of normative functional neuroimaging studies. Human Brain Mapping, 25(1), 46–59. https://doi.org/10.1002/hbm.20131

Puts, N. A. J., Edden, R. A. E., Evans, C. J., McGlone, F., & McGonigle, D. J. (2011). Regionally Specific Human GABA Concentration Correlates with Tactile Discrimination Thresholds. The Journal of Neuroscience, 31(46), 16556–16560. https://doi.org/10.1523/JNEUROSCI.4489-11.2011

Sandberg, K., Blicher, J. U., Dong, M. Y., Rees, G., Near, J., & Kanai, R. (2014). Occipital GABA correlates with cognitive failures in daily life. NeuroImage, 87, 55–60. https://doi.org/10.1016/j.neuroimage.2013.10.059

Shibata, K., Sasaki, Y., Bang, J. W., Walsh, E. G., Machizawa, M. G., Tamaki, M., … Watanabe, T. (2017). Overlearning hyperstabilizes a skill by rapidly making neurochemical processing inhibitory-dominant. Nature Neuroscience. https://doi.org/10.1038/nn.4490

Song, C., Sandberg, K., Andersen, L. M., Blicher, J. U., & Rees, G. (2017). Human Occipital and Parietal GABA Selectively Influence Visual Perception of Orientation and Size. Journal of Neuroscience, 37(37), 8929–8937. https://doi.org/10.1523/JNEUROSCI.3945-16.2017

van Loon, A. M., Knapen, T., Scholte, H. S., St. John-Saaltink, E., Donner, T. H., & Lamme, V. A. F. (2013). GABA Shapes the Dynamics of Bistable Perception. Current Biology, 23(9), 823–827. https://doi.org/10.1016/j.cub.2013.03.067

Waddell, K. W., Avison, M. J., Joers, J. M., & Gore, J. C. (2007). A Practical Guide to Robust Detection of GABA in Human Brain by J-difference Spectroscopy at 3 Tesla Using a Standard Volume Coil. Magnetic Resonance Imaging, 25(7), 1032. https://doi.org/10.1016/j.mri.2006.11.026

Wagenmakers, E.-J., Love, J., Marsman, M., Jamil, T., Ly, A., Verhagen, J., … Morey, R. D. (2017). Bayesian inference for psychology. Part II: Example applications with JASP. Psychonomic Bulletin & Review, 1–19. https://doi.org/10.3758/s13423-017-1323-7

Wagenmakers, E.-J., Marsman, M., Jamil, T., Ly, A., Verhagen, J., Love, J., … Morey, R. D. (2017). Bayesian inference for psychology. Part I: Theoretical advantages and practical ramifications. Psychonomic Bulletin & Review, 1–23. https://doi.org/10.3758/s13423-017-1343-3

Yoon, J. H., Grandelis, A., & Maddock, R. J. (2016). Dorsolateral Prefrontal Cortex GABA Concentration in Humans Predicts Working Memory Load Processing Capacity. Journal of Neuroscience, 36(46), 11788–11794. https://doi.org/10.1523/JNEUROSCI.1970-16.2016

Zhang, J., & Mueller, S. T. (2005). A note on ROC analysis and non-parametric estimate of sensitivity. Psychometrika, 70(1), 203–212. https://doi.org/10.1007/s11336-003-1119-8

